# Response to Qian et al (2017): Daily and seasonal climate variations are both critical in the evolution of species’ elevational range size

**DOI:** 10.1101/195248

**Authors:** Wei-Ping Chan, I-Ching Chen, Robert K. Colwell, Wei-Chung Liu, Cho-ying Huang, Sheng-Feng Shen

## Abstract

In their recent critique, Qian et al. (2017) claimed that the results of structural equation modeling analysis (SEM) in Chan et al. (2016) were flawed. Here, we show that the source of the difference in their re-analysis is that Qian et al. did not follow the standard, iterative process of SEM, which allows researchers to evaluate which model offers the best account of the data in both absolute and relative senses. Here, we provide step-by-step instructions to reproduce our published results. All of Qian et al.’s concerns regarding SEM can be put to rest. Moreover, in our original paper we used three distinct statistical methods—hierarchical partitioning, SEM, and stationary bootstrap—to show that different temporal scales of environmental variability can differentially impact the elevational range size (ERS) of species. It is time to move on to probing the pressing issue of how and why climatic variability impacts ERS.

---

In a critique of Chan *et al*. (2016), Qian et al. (2017) concluded that the structural equation modeling (SEM) results in our Figure 1a (Chan *et al.*, 2016) were flawed, because they could not repeat our results, “using the same dataset, analysis, and software.” In fact, it seems that they were unable to repeat most of our SEM results, stating, “…we cannot replicate the results reported for model 3 in Chan et al. (2016), nor those for many of the other models reported in their Table S2”. A close reading of Qian et al.’s Supporting Information explains how they arrived this conclusion. Evidently, they did not follow the standard, iterative process that distinguishes SEM from traditional statistical tests, consisting of five consecutive (and iterative) steps: (1) model specification, (2) model identification, (3) estimation, (4) evaluation of fit, and (5) model modification (Chou & Huh, 2012; Hoyle, 2012a, b). Thus, it is understandable that Qian et al. did not obtain the same results we did. In retrospect, we regret not having been more comprehensive in specifying the details of our SEM analyses in the Supplemental Online Material of Chan et al. (2016).

**Figure 1.**
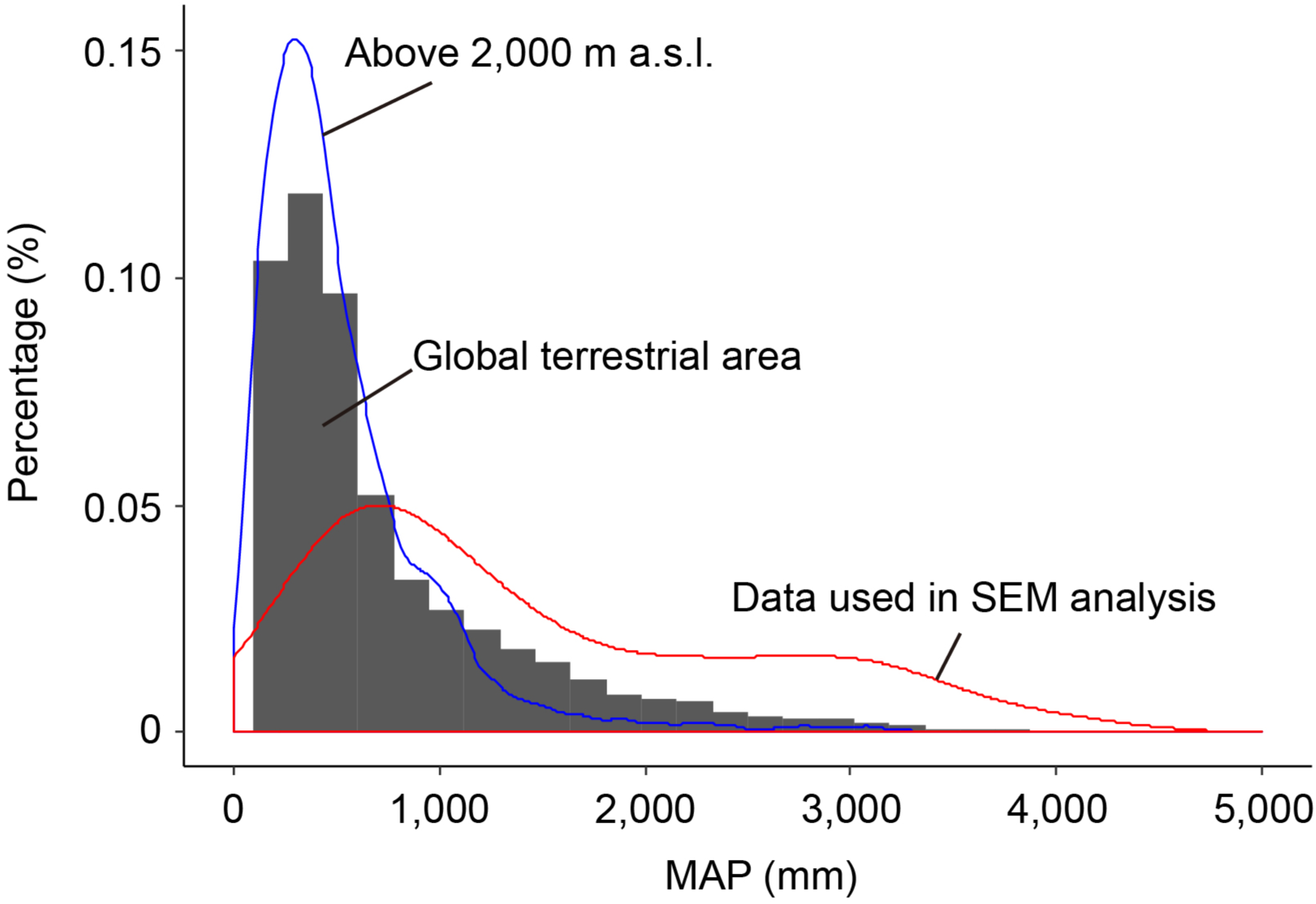
The global frequency distributions of mean terrestrial annual precipitation (MAP) and land above 2,000m, compared to the data used in our SEM analysis. The distribution of the MAP data in our SEM analysis was significantly wetter than the global distribution of MAP (Wilcoxon signed-rank test; w = 2 071 600, p < 0.0001), and it was also significantly wetter than the area of land above 2,000 m (Wilcoxon signed-rank test; w = 74 338, p < 0.0001).

To clarify every step of every procedure conducted in our analysis, we have prepared additional Supplemental Materials for Chan et al. (2016) [submitted to *Science*, for consideration as additional Supplemental Material for our published paper (Chan *et al*. 2016), but provided for editors and reviewers of this submission to *J. Biogeogrpaphy*], which provide full instructions and materials for repeating and refining the statistical results reported in Chan et al. (2016), including the raw and intermediate data, all source code, and the table of parameters for reproducing the results that were reported in Chan et al. 2016 (data can be downloaded in https://data.mendeley.com/datasets/mkd68skn5z/2). By following our instructions, we expect that anyone with access to the appropriate software (IBM SPSS AMOS, (Arbuckle, 2012)) can easily apply our approach and reproduce our results. All of Qian et al.’s concerns regarding the reported coefficients and criteria for model-fit can be put to rest. (We are puzzled by Qian et al.’s claim that they used the same software we did. While we used IBM SPSS AMOS, Qian et al. apparently used LAVAAN in R, although we believe the correct use of different packages should give the similar results.)

Qian et al.’s neglect of the iterative nature of SEM analysis perhaps reflects a deeper misunderstanding of SEM analysis. Many researchers, including ecologists, have repeatedly emphasized the hypothesis-testing nature of SEM analysis (Grace, 2006; Chou & Huh, 2012; Hoyle, 2012a). Thus, we build models to represent a number of theoretically-meaningful, candidate hypotheses, based on the literature and our subjective judgment, and subsequently, use objective statistical criteria for model-fit to develop one or more statistically adequate models. There are two ways to evaluate the alternative models. In an absolute sense, we evaluate whether each model has a reasonable fit to the observed data; and, in a relative sense, we evaluate which model offers the best account of the data.

Mathematically, an SEM is represented by a set of simultaneous equations. If there are too many free variables (parameters, coefficients, and/or intercepts) in a candidate model to produce an adequate solution of these simultaneous equations, or the data are not sufficient to support a statistically adequate solution, the value of one or more of the variables can be constrained (specified). Thus, when we found that the fit of a candidate model was inadequate (by assessing the ratio of chi-squared to degrees of freedom, which must be no greater than 5 (Jöreskog, 1969; Wheaton *et al.*, 1977; West *et al.*, 2012)), we simplified the model (i.e. model modification), where possible and justifiable, by decreasing the number of free variables (parameters, path coefficients, and/or intercepts) to be evaluated in the SEM analysis by specifying the value of one or more variables, thereby increasing the degrees of freedom of the model. In SEM, specifying the value of a variable in a model, instead of estimating it from the model, is a means of *identifying* the variable.

All the alternative model structures that we proposed (Table S2 in Chan et al. 2016) required simplification by the specification of variables to achieve acceptable degrees of freedom, under the chi-squared ratio criterion. These simplifications are fully detailed in the Additional Supplemental Material [submitted to Science for online publication], in which we describe additional technical details and reproduce in detail the key results in Chan *et al.* 2016. To fairly compare different alternative models that required specification of variables, we first explored the influences of different model identification strategies on model fit, for each model. Specifically, we compared models with (1) different numbers and (2) different positions (i.e. primary or secondary hierarchy) of identified (specified) parameters and path coefficients. We found that model fit values varied very little among different identification strategies (see Table S8 and S9). In all cases of acceptable models (RMSEA<0.08 and CFI >0.95), SRMR ranged only between 0.0218 and 0.0243 for different numbers of identifications (Table S8) and between 0.0217 to 0.0241 for different positions (Table S9), respectively). Thus, we chose (in Chan et al. 2016) to uniformly identify parameters and path coefficients to represent our results. Specifically, we specified the means and/or variances for a uniform set of parameters and path coefficients (the means of Latitude = 0.497, MAP = 0.322, and DTR = 0.541; the multiple regression coefficients between Latitude and STR = 0.830 and between DTR and range size = −0.233). These values can be easily computed from the model input data using any statistical tool (e.g. SPSS or R).

By this means, we were able to decrease number of free parameters and appropriately modify our models, guided by statistical evaluations of fit, and determine which candidate model was the best model, as we reported in Chan et al. (2016). In contrast, without employing the steps of model specification, identification, estimation, evaluation of fit, and model modification required for the proper and most effective use of SEM analysis, it is perhaps not surprising that Qian et al. concluded, “We are unable to meaningfully improve on the analysis of this dataset that was published by McCain (McCain, 2009), so we do not attempt to provide a new ‘best model’.” In short, we used SEM to evaluate alternative hypotheses, which helped us identify a better candidate hypothesis. We then performed further analyses, using stationary bootstrap, to validate the nature of our SEM findings. We found that mean annual precipitation (MAP) plays a more important role in influencing the relative importance of DTR and STR in affecting the elevational range size of species.

Many of Qian et al.’s (2017) other concerns have been adequately addressed in our original paper (Chan *et al.*, 2016), including the most appropriate and powerful statistical approaches to deal with uneven or pseudoreplicated sampling arising from aggregated geographic distributions or evolutionary relatedness of taxonomic groups. These are common issues in macroecological studies, and conventionally, rarefying samples or dichotomizing continuous predictors are widely-adopted procedures, without considering the serious statistical and inferential perils of these approaches. Instead, we conducted formal statistical analyses to deal with these sampling issues that use all the data and do not depend on arbitrary dichotomies.

Our analysis shows that latitude and MAP emerged as the geographical and environmental factors that indirectly shape elevational range size, through their influence on climatic variability (diurnal temperature range or DTR, and seasonal temperature range, or STR). As Qian et al. correctly point out, mountain gradients in the McCain (2009) dataset are not evenly distributed across Earth’s continents. It is precisely for this reason that we used a stationary bootstrap method to assess whether STR and/or DTR is more explanatory than expected at random along gradients of latitude and mean annual precipitation (MAP, representing the wet-arid continuum). The issue of uneven sample distribution is commonly faced by most, if not all, meta-analysis studies. Stationary bootstrap—a block-resampling procedure that joins blocks of random length—is a formal statistical method for dealing with uneven sampling along a stationary gradient, such as a time series, latitude, or an environmental variable, such as precipitation (Politis & Romano, 1992, 1994). Using stationary bootstrap analysis, we showed that DTR is more important than STR in explaining elevational range size (ERS) in regions with mid-MAP (1000mm-1600mm) and in mid-latitude regions (25-35 N), whereas STR is more important than DTR in regions with MAP between 1600mm and 2700m, but not in any particular latitude (Fig. 2 of Chan et al., 2016).

It might seem appropriate to rarefy the over-represented dry mountain moiety of the dataset to make the proportion of samples fit better to the proportion of dry and wet gradients in real world, as Qian et al. did by removing half of the data from the drier regions. Clearly, it is tempting to set up a qualitative dichotomy, i.e. between “wet” and “dry” conditions, for a continuous variable and evaluate results separately (as done by Qian et al.). However, the perils of dichotomizing continuous predictors—including the arbitrariness of the cut-off point and inevitably smaller sample sizes—are serious, and have been extensively discussed (Royston *et al.*, 2006). Thus, separating the data into “arid” vs. “wet” gradients and performing the SEM separately for each group of gradients is statistically weak and potentially misleading. In fact, if we compare the distribution of climatic conditions, by surface area, globally, low-rainfall regions are actually under-represented in our analysis (Figure 1). Thus, the role of DTR in influencing elevational range size of species might be even more important than suggested by our analysis. Overall, applying the stationary bootstrap procedure is a more repeatable, non-arbitary, and rigorous approach to deal with this issue, although the method is not familiar to most macroecologists. We suggest that macroecologists follow our lead, and move away from arbitrarily removing data and setting up an arbitrary dichotomies, instead using more a more formal resampling method on a continuous variable to resolve the issue of the uneven sample distributions. We are not persuaded by the argument that any particular statistical procedure should be followed simply because it is familiar and commonly used, once manifestly better approaches have been developed.

Qian et al. criticized our decision not to report *r*^2^ in our SEM analysis. Instead, we provided a more comprehensive analysis of *r*^2^ for the effects of DTR and STR in our stationary bootstrap analysis (in our original Fig. 2A), with the comparison of observed to random expectation of r^2^. This approach is more rigorous than the approach that Qian et al. used in their critique.

Qian *et al*. argued against taxonomically comprehensive analysis, adding that “McCain (pers. comm.) strongly cautions against this, arguing in particular that including the rodent data in the analysis is inappropriate because rodents have the opposite elevational range size trend to the other vertebrate groups….[O]ur final reanalysis started by excluding the rodent data and shows that removing a single, pseudoreplicated data-point again overturns the key empirical conclusion of Chan et al. (2016).” In addition to taxonomic sub-setting, we note that McCain’s (2009) key analyses were based on simple regression, with elevational data truncation (elevational distribution data were truncated at certain, taxon-specific elevations and only range sizes below these elevation were compared across latitude in a series of regressions), without considering the inherent collinearity among candidate predictors and the pitfalls of data sub-setting. In our paper, we formally examined the taxon-specific effect on range sizes (Fig. S11 in Chan *et al.* 2016). We found that DTR has a stronger effect on rodents, snakes and lizards than on other taxa. There is no *a priori* justification for removing any of these groups from the dataset. In fact, in an alternative approach, we also statistically controlled for taxonomic effect by setting taxon as a variable in the model, as described in the Supplementary Material for our published paper (Chan et al. 2016, p.3, L91-92). Following McCain’s approach (McCain, 2009), Qian et al. also used simple regression to conclude that, “the diurnal temperature range and mean elevational range size variables are not correlated with each other (r = .039, p = .651; Figure 1a).” Simple regressions on 2-variable scatterplots can completely miss important and significant relationships, when confounding variables are not included in the analysis. It is straightforward to show that, if DTR and STR are both included in a multiple regression model, both DTR and STR have significant effects on the elevational range size of species (for DTR: t= −2.19, p = 0.03 for STR: t = 3.72, p < 0.01).

For readers interested in whether daily climate variation impacts the elevational range size (ERS) of species, we remain confident that the answer is, “Yes”. We have used three statistical methods—hierarchical partitioning, structural equation modelling (SEM), and stationary bootstrap—to show that daily temperature range (DTR) has a significantly negative influence on ERS, and that its explanatory power is, in general, higher than seasonal temperature range (STR), a factor widely viewed as representing climatic variation (e.g. DTR explains 13.8%, whereas STR explains 2.9% of explainable variation in ERS; Supplementary Material of Chan et al. 2016).

Following Janzen’s insightful climatic variability hypothesis (Janzen, 1967) and Gilchrist’s simple but elegant model (Gilchrist, 1995), we offered evidence for the hypothesis that different temporal scales of environmental variability can differentially impact the ERS of species. Many interesting puzzles regarding the global pattern of ERS emerge from our study, such as whether endotherms and ectotherms respond differently to DTR and STR. By what mechanisms does larger DTR promote niche specialization, and, thus, smaller range size of species (other things being equal)? And, if climate change poses a greater threat to species with smaller range sizes, as many believe (e.g. Tewksbury *et al.*, 2008; Perez *et al.*, 2016), will species in regions with large DTR be more vulnerable, just as tropical species in regions with lower STR are believed to be especially vulnerable? More studies on the impacts of climatic variability on species range size are clearly needed.

